# FUT8-mediated core fucosylation modulates growth-related functions of LRP1 in liver cancer cells

**DOI:** 10.64898/2026.04.09.717446

**Authors:** Asuka Ogata, Mika Ueda, Koki Ohyama, Shinji Takamatsu, Yuki Makino, Hayato Hikita, Yoshiyuki Manabe, Koichi Fukase, Yusuke Oji, Yoshihiro Kamada, Jumpei Kondo, Eiji Miyoshi

**Affiliations:** 1 Department of Molecular Biochemistry and Clinical Investigation, Graduate School of Medicine, The University of Osaka, Suita, Osaka, Japan; 2 Department of Gastroenterology and Hepatology, Graduate School of Medicine, The University of Osaka, Suita, Osaka, Japan; 3 Department of Chemistry, Graduate School of Science, The University of Osaka, Toyonaka, Osaka, Japan; 4 Department of Clinical Laboratory and Biomedical Sciences, Graduate School of Medicine, The University of Osaka, Suita, Osaka, Japan; 5 Department of Advanced Metabolic Hepatology, Graduate School of Medicine, The University of Osaka, Suita, Osaka, Japan

## Abstract

Fucosylation is a cancer-associated glycosylation change, and core fucosylation of N-glycans catalyzed by the α1,6-fucosyltransferase FUT8, has been closely linked to tumor progression, metastasis, drug resistance, and poor prognosis. However, the core-fucosylated proteins that directly support hepatocarcinogenesis are not fully defined. In a KRAS-G12D-driven mouse liver cancer model, we observed increased fucosylation and FUT8 upregulation, and glycoproteomic analysis of fucose-enriched fractions identified low-density lipoprotein receptor-related protein 1 (LRP1) as a prominent core-fucosylated protein in tumor tissue. In immortalized mouse hepatocytes, genetic or pharmacological inhibition of FUT8 markedly increased *Lrp1* mRNA and protein, indicating that loss of core fucosylation is accompanied by robust upregulation of LRP1. In human HepG2 cells, LRP1 knockout suppressed cell proliferation and markedly altered colony morphology, leading to compact rounded clusters instead of the typical polygonal pattern. It also reduced EGFR protein and further inhibited proliferation of HepG2 cell. These findings identify LRP1 as a FUT8-dependent core-fucosylated receptor in experimental hepatocarcinogenesis and suggest that the FUT8–LRP1 axis contributes to the maintenance of proliferative signaling in hepatoma cells.

## Introduction

Hepatocellular carcinoma (HCC) is one of the leading causes of cancer-related death worldwide, and its incidence continues to rise despite advances in antiviral therapy and surveillance. Aberrant protein glycosylation is a hallmark of malignant transformation, and changes in *N*-glycan structures have been repeatedly documented in HCC(1–3). Among these alterations, increased fucosylation, particularly core fucosylation of *N*-glycans catalyzed by the α1,6-fucosyltransferase FUT8, has been closely linked to tumor progression, metastasis, drug resistance, and poor prognosis(2, 4, 5). Fucosylated glycoproteins such as α-fetoprotein-L3(6, 7) and fucosylated haptoglobin(8–10) have already been proposed as biomarkers for liver cancer, yet the specific core-fucosylated molecules that functionally contribute to hepatocarcinogenesis remain incompletely defined.

FUT8 is the sole enzyme responsible for adding core fucose to complex and hybrid *N*-glycans in mammals(11, 12), and *Fut8*-deficient mice exhibit profound developmental and pathological abnormalities(13), underscoring the essential role of core fucosylation in receptor signaling and tissue homeostasis(14). In the liver, comprehensive *N*-glycan profiling and tissue studies have shown that FUT8 expression and core-fucosylated *N*-glycans are increased in HCC and correlate with malignant behavior of HCC cells(1, 4). Moreover, FUT8-mediated core fucosylation modulates the activity of several key receptors, including TGF-β(14, 15) and EGF receptors(16), thereby influencing proliferation, epithelial–mesenchymal transition, and multidrug resistance in cancer. Despite these advances, only a limited number of FUT8 substrates have been functionally characterized in hepatocytes and hepatoma cells, which hampers a detailed understanding of how FUT8 links altered glycosylation to oncogenic signaling in HCC.

Beyond its role as a biomarker-related enzyme, FUT8 has therefore emerged as a potential therapeutic target, particularly in the context of cancer immunotherapy and pathway-directed therapies. Antibody-producing cells with reduced FUT8 expression generate defucosylated therapeutic antibodies with enhanced antibody-dependent cellular cytotoxicity (ADCC)(17, 18). Several studies suggest that inhibiting FUT8 in immune cells can reprogram immune receptor signaling and immune checkpoint molecules such as PD-1 and B7H3 to improve antitumor responses(19, 20). In parallel, small-molecule and mechanism-based FUT8 inhibitors have been developed(20, 21), including recent compounds that selectively inhibit FUT8 in cells(22, 23). These advances demonstrate the feasibility of pharmacologically targeting core fucosylation in living systems. However, in HCC, it remains largely unclear which core-fucosylated receptors and co-receptors in hepatocytes are critical for sustaining oncogenic signaling, or which of them might mediate unwanted effects of FUT8 inhibition, making it essential to define functionally relevant FUT8 substrates before FUT8-targeted interventions can be rationally evaluated in this disease.

Here, we sought to clarify how FUT8-dependent core fucosylation contributes to pathophysiology of HCC by identifying core-fucosylated glycoproteins in mouse liver tumor. Using lectin-based glycoproteomic analysis, we identified Low density lipoprotein receptor-related protein 1 (LRP1) as a core-fucosylated glycoprotein associated with FUT8 upregulation, and then examined how genetic or pharmacological inhibition of FUT8 affects its expression in hepatocytes. We further assessed the functional role of LRP1 in the human hepatoma cell line HepG2, focusing on its impact on cell proliferation.

## Materials and Methods

### Mice

All animal experiments were approved by the Animal Ethics Committee of the Department of Health Sciences, The University of Osaka Graduate School of Medicine and were performed in accordance with institutional guidelines.

For the genetically engineered model, C57BL/6/129 background mice carrying a loxP-stop-loxP (LSL) cassette followed by a *Kras^G12D^* allele (*Kras^LSL-G12D/+^*) were kindly provided by the National Cancer Institute (Bethesda, MD, USA). Hepatocyte-specific Kras-mutant mice (*Kras^LSL-G12D/+^*;*Alb-Cre*) were generated by crossing *Kras^LSL-G12D/+^* mice with heterozygous *Alb-Cre* transgenic mice expressing Cre recombinase under the control of the mouse albumin promoter (24). Mice were sacrificed at 8–11 months of age, and liver tissues were collected from macroscopically cancerous lesions and from regions without any grossly apparent tumors in the same livers.

For the chemical carcinogenesis model, male C57BL/6J wild-type mice received a single intraperitoneal injection of diethylnitrosamine (DEN, Sigma-Aldrich Japan, Tokyo) at 2 weeks of age as described previously(25). Mice were euthanized at 9 months of age, and liver samples were obtained from visible cancerous nodules and from liver regions without grossly detectable tumors. Normal liver tissue was collected from age-matched male C57BL/6J wild-type mice without any treatment.

### Cell culture

All cells were maintained in Dulbecco’s modified Eagle’s medium (DMEM; Nacalai Tesque, Kyoto, Japan) supplemented with 10% fetal bovine serum (FBS; NICHIREI BIOSCIENCE, Tokyo, Japan), 100 U/mL penicillin, and 100 μg/mL streptomycin (Nacalai Tesque) in a humidified incubator at 37°C with 5% CO₂. An immortalized mouse hepatocyte cell line derived from wild-type and Fut8-knockout mice (LivSV40 WT and LivSV40 KO) was established previously (26). HepG2 cells were obtained from the American Type Culture Collection (ATCC, Manassas, VA, USA). HepG2 LRP1-knockout cells were generated using the lentiviral vector pLenti-CRISPR v2 (Addgene, Watertown, MA, USA), which expresses Cas9 and a guide RNA targeting LRP1. The guide RNA sequence was 5’-GGGCCTCGTCAGATCCGTCT-3’.

For chemical inhibition of FUT8 and general fucosylation in LivSV40 WT cells, a selective FUT8 inhibitor (Compound 40) (22), and a commercially available fucosylation inhibitor, 2F-peracetylfucose (Merck, Rahway, NJ, USA), were used at final concentrations of 2 μM and 100 μM, respectively. Because of its poor stability in culture medium, the FUT8 inhibitor was added repeatedly at 0, 6, 24, 30, 48, 54, and 72 hours after seeding. Cells were harvested at 78 hours for western blot analysis.

Cell proliferation was assessed using the CellTiter-Glo® (Promega, Madison, WI, USA). Cells were seeded into 96-well plates at 1000 cells/well and cultured for 1, 3, and 5 days, after which CellTiter-Glo reagent was added according to the manufacturer’s instructions. Luminescence was measured using an Infinite M plex plate reader (model 200 PRO, manufacturer TECAN).

### Lectin blot and western blot

Equal amounts of protein from tissue extracts or cell lysates were separated by SDS-PAGE and transferred onto PVDF membranes. For lectin blotting, fucosylated proteins were detected using biotin-labeled Aleuria aurantia lectin (AAL; J-Oil Mills, Tokyo, Japan) followed by streptavidin–HRP (ab7403; Abcam, Cambridge, UK).

For western blotting, membranes were first incubated with primary antibodies and then with HRP-conjugated secondary antibodies as listed in Supplementary Table S1, followed by detection with an ECL chemiluminescence reagent (GE Healthcare, Chicago, IL, USA). Chemiluminescent signals were visualized using a ChemiDoc Touch imaging system (Bio-Rad Laboratories, Hercules, CA, USA).

### Lectin Precipitation

Extracted proteins (100 μg) were incubated with 1 μL of biotin-labeled AAL overnight at 4°C and then mixed with pre-cleaned streptavidin Sepharose (40 μL, 1:1 slurry; Cytiva, Uppsala, Sweden) for 1 hour at 4°C. The precipitated proteins were washed, eluted by boiling in SDS sample buffer, and separated by SDS-PAGE. Proteins were visualized by silver staining using the Silver Stain MS Kit (FUJIFILM Wako, Osaka, Japan) according to the manufacturer’s instructions. Mass spectrometry analysis of silver-stained bands of interest was performed by the Joint Research Center for Medical Research and Education at The University of Osaka (Osaka, Japan).

### Semi-quantitative real time RT-PCR

Total RNA was extracted from cells using the RNeasy Mini Kit (Qiagen, Venlo, Netherlands). One microgram of RNA was reverse transcribed into complementary DNA using the ReverTra Ace qPCR RT Kit (TOYOBO, Osaka, Japan). Quantitative real-time PCR was performed with THUNDERBIRD SYBR qPCR Mix (TOYOBO) on a QuantStudio™ real-time PCR system (Life Technologies, Carlsbad, CA, USA). Gene expression levels were calculated using the comparative Ct (ΔΔCt) method with Gapdh as an internal reference. The primer sequences were as follows: *Lrp1* Fwd; 5’-GAACCACCATCGTGGAAA-3’, Rev; 5’TCCCAGCCACGGTGATAG-3’, *Gapdh* Fwd; 5’-TGGTGCTGCCAAGGCTGTGG-3’, REV; 5’-GGCAGGTTTCTCCAGGCGGC-3’

### Flow cytometry

HepG2 wild-type and LRP1-knockout cells were seeded at 5.0 × 10⁵ and 1.0 × 10⁶ cells per well, respectively, in 6-well plates and cultured for 3 days. Cells were detached with trypsin/EDTA solution (Nacalai Tesque), fixed in cold 70% ethanol (FUJIFILM Wako) overnight at 4°C, washed, treated with RNase (Roche), and stained with 5 μg/mL propidium iodide (Nacalai Tesque). Cell-cycle distribution was analyzed using an Attune NxT Acoustic Focusing Cytometer (Thermo Fisher Scientific).

### Statistical Analysis

For comparisons between two groups, a two-tailed Student’s t-test was used. For multiple comparisons, Student’s t-tests with Bonferroni correction were applied. A p value < 0.05 was considered statistically significant.

## Results

### Increased core fucosylation and FUT8 expression in Kras-driven, but not DEN-induced, liver tumors

To examine changes in fucosylation during hepatocarcinogenesis, we first analyzed two mouse liver cancer models. In hepatocyte-specific *Kras^LSL-G12D/+^*; *Alb-Cre* mice, AAL lectin blotting revealed a clear increase in fucosylated proteins in tumor tissue compared with liver regions without grossly detectable tumors (hereafter, non-tumor liver) and normal liver (Fig. 1A). In the same Kras-driven tumors, FUT8 protein was markedly upregulated by western blotting (Fig. 1B). In contrast, neither AAL signals nor FUT8 expression were appreciably increased in DEN-induced liver tumors compared with non-tumor liver in the same animals (Fig. 1A, B). These results indicate that core fucosylation and FUT8 are selectively enhanced in the Kras-driven HCC model.

**Figure 1.**
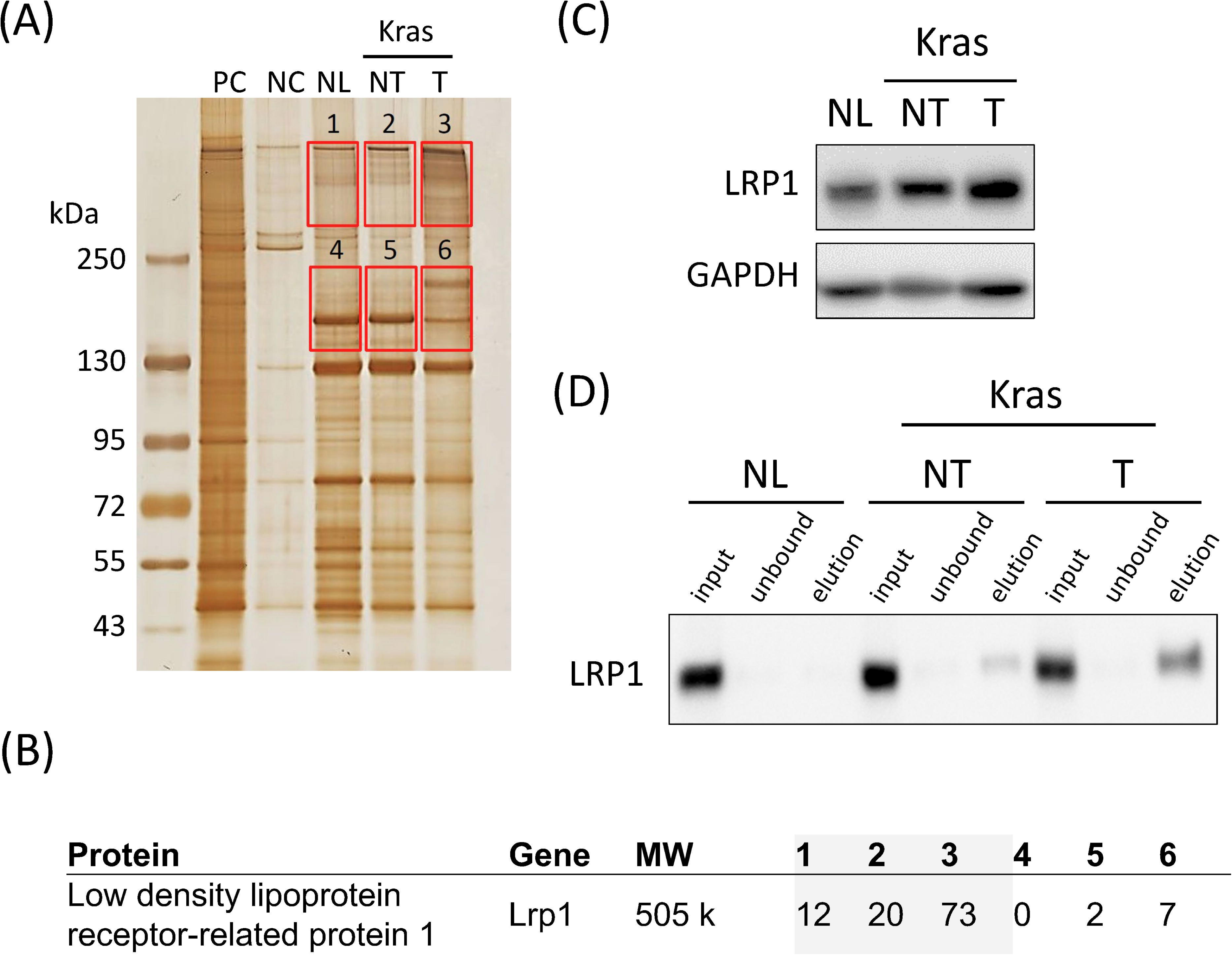
Increased core fucosylation and FUT8 expression in Kras-driven, but not DEN-induced, liver tumors (A) AAL lectin blot of protein lysates from human pancreatic ductal adenocarcinoma PK45 cells (positive control, PC), human colorectal cancer HCT116 cells lacking fucosylated glycans (negative control, NC), normal liver tissue from untreated mice (NL), and paired non-tumor liver (NT) and tumor tissues (T) from DEN-induced (DEN) and Kras-driven tumor–bearing mice (Kras). (B) Western blot analysis of FUT8 in NL and in paired NT and T samples from DEN and Kras mice; GAPDH was used as a loading control.

### Identification of LRP1 as an aberrantly fucosylated protein in Kras mouse liver tumor

To identify tumor-associated fucosylated proteins, we performed AAL lectin precipitation followed by SDS-PAGE and silver staining of liver lysates from *Kras^LSL-G12D/+^*; *Alb-Cre* mice. A tumor-specific band with increased intensity was observed in AAL-precipitated fractions from cancerous liver tissue than normal liver or liver regions without grossly detectable tumors of Kras mice (Fig. 2A). Mass spectrometry analysis of this band identified LDL receptor-related protein 1 (LRP1) as a major component (Fig. 2B, Supplementary Table. S2). Western blotting showed that total LRP1 protein levels were higher in Kras tumor tissue than in normal liver or non-tumor liver from Kras mice (Fig. 2C). Furthermore, when comparable amounts of LRP1-containing input were applied to an AAL affinity column, larger amounts of AAL-reactive LRP1 were recovered in the eluted fractions from Kras tumor lysates than from normal or non-tumor liver (Fig. 2D). These findings indicate that LRP1 expression and its fucosylation are both increased in Kras-driven mouse liver tumors.

**Figure 2.**
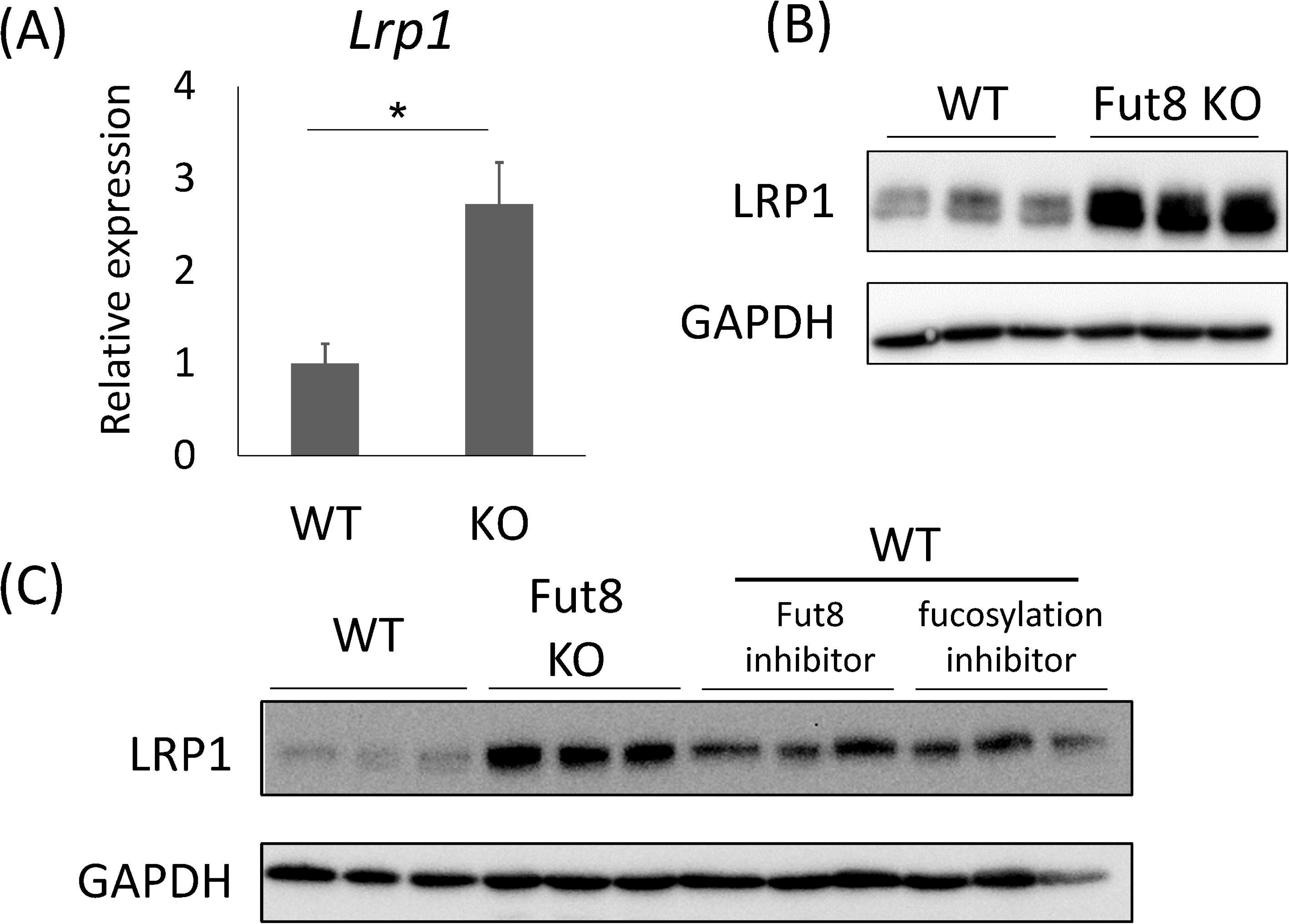
Identification of LRP1 as an aberrantly fucosylated protein in Kras mouse liver tumors (A) Silver-stained SDS-PAGE of proteins precipitated with AAL from LivSV40 wild-type cells (positive control, PC), LivSV40 Fut8-knockout cells (negative control, NC), normal liver tissue from wild-type mice (NL), and non-tumor liver (NT) and tumor (T) tissues from Kras-driven tumor–bearing mice. The region enclosed by the red box was used for MS proteomics analysis. The area numbers correspond to those in Fig. 2B and Supplementary Table S2. (B) For LRP1, the spectrum counts corresponding to each lane in Fig. 2A are shown. (C) Western blot analysis of LRP1 in NL, NT, and T tissues from Kras mice; GAPDH was used as a loading control. (D) AAL affinity chromatography of liver lysates from wild-type normal liver (NL), and NT and T tissues from Kras mice using input samples adjusted to obtain comparable LRP1 levels. LRP1 in the input, unbound flow-through, and eluted fractions was analyzed by western blotting.

### FUT8 deficiency or inhibition upregulates LRP1 in immortalized mouse hepatocytes

We next examined how FUT8 activity affects LRP1 expression in hepatocytes using immortalized mouse hepatocyte lines (LivSV40) derived from wild-type and Fut8-knockout mice. In LivSV40 cells, Fut8 knockout increased *Lrp1* mRNA levels by approximately 2.7-fold compared with wild-type cells (Fig. 3A) and caused a striking accumulation of LRP1 protein (Fig. 3B). AAL lectin precipitation further confirmed that total LRP1 protein was elevated in Fut8-knockout cells despite the loss of core fucosylation (Fig. 3B). In wild-type LivSV40 cells, treatment with the selective FUT8 inhibitor Compound 40 or the global fucosylation inhibitor 2F-peracetylfucose also led to increased LRP1 protein expression (Fig. 3C). These results demonstrate that both genetic and pharmacological inhibition of FUT8 are accompanied by robust upregulation of LRP1 in hepatocytes.

**Figure 3.**
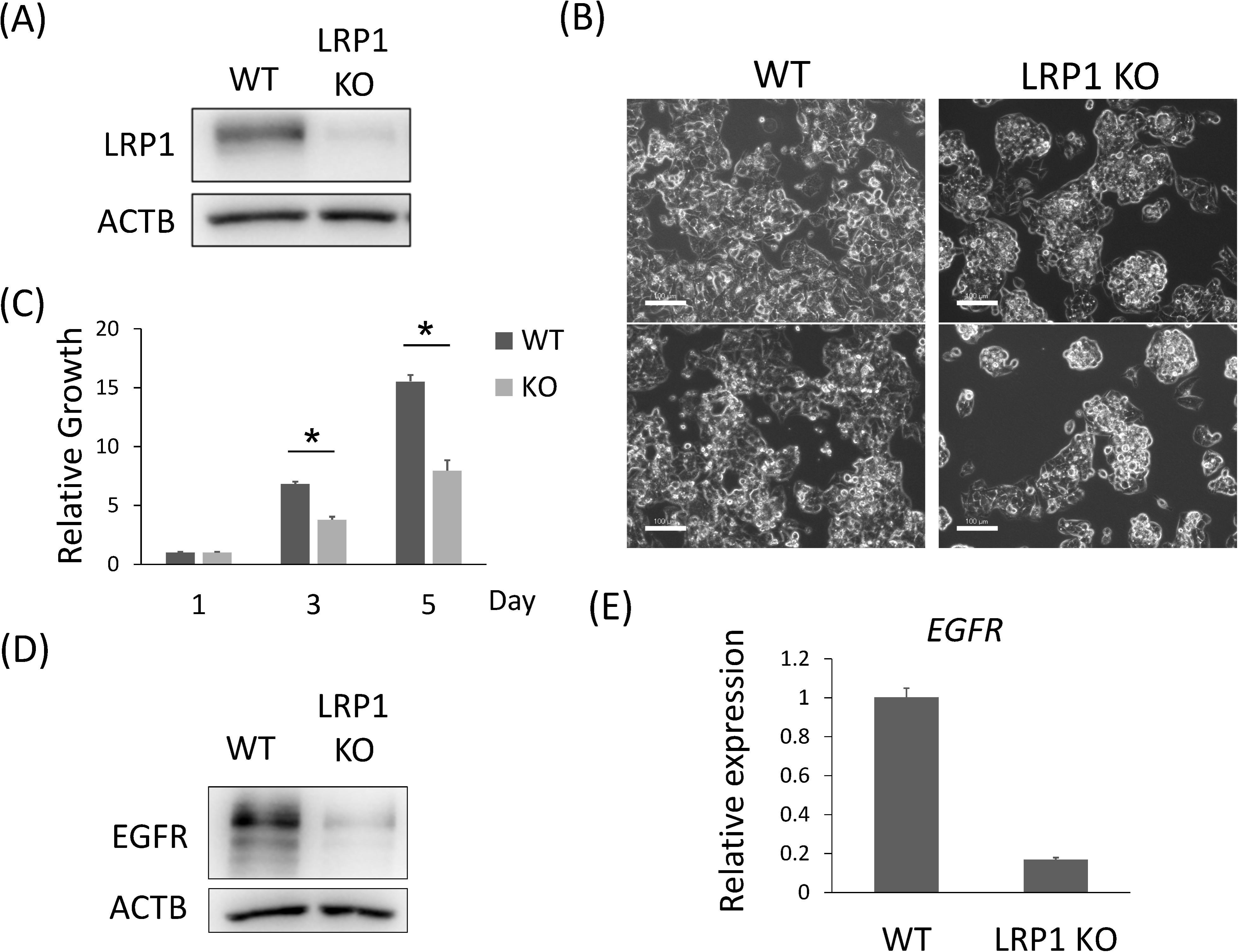
FUT8 deficiency or inhibition upregulates LRP1 in immortalized mouse hepatocytes (A) Semi-quantitative RT-PCR analysis of *Lrp1* mRNA expression in LivSV40 wild-type (WT) and LivSV40 Fut8-knockout (KO) cells. Expression levels were normalized to *Gapdh* and are shown relative to WT cells (set to 1.0); n = 3 per group, mean ± SE, *p < 0.05 (Student’s t-test). (B) Western blot analysis of LRP1 protein in LivSV40 WT and Fut8 KO cells (three independent samples per group); GAPDH was used as a loading control. (C) Western blot analysis of LRP1 in LivSV40 WT cells treated with a selective FUT8 inhibitor (compound 40) or a pan-fucosylation inhibitor (2F-peracetylfucose), and in Fut8 KO cells (three independent samples per group); GAPDH was used as a loading control.

### LRP1 deficiency suppresses proliferation and downregulates EGFR expression in HepG2 cells

To clarify the functional role of LRP1 in hepatoma cells, we generated LRP1-knockout HepG2 cells using CRISPR/Cas9. After antibiotic selection, only a single resistant clone survived, and western blotting confirmed the loss of LRP1 protein in this clone (Fig. 4A). Phase-contrast microscopy revealed a marked change in morphology: whereas wild-type HepG2 cells formed a typical spreading monolayer with polygonal morphology, LRP1-knockout cells grew as compact, rounded clusters (Fig. 4B). In cell proliferation assays, LRP1-knockout HepG2 cells grew significantly more slowly than wild-type cells over 1, 3, and 5 days in culture (Fig. 4C). Western blot analysis showed that total EGFR protein levels were decreased in LRP1-knockout HepG2 cells compared with wild-type cells (Fig. 4D), and quantitative RT-PCR confirmed a concomitant reduction in *EGFR* mRNA expression (Fig. 4E). In the LRP1-knockout clone, flow-cytometric analysis of propidium iodide-stained cells revealed an increased proportion of cells in the G2/M phase compared with wild-type cells (Supplementary Fig. S1). These data indicate that LRP1 deficiency is associated with reduced EGFR abundance and altered cell-cycle distribution, consistent with reduced proliferation in HepG2 cells.

**Figure 4.**
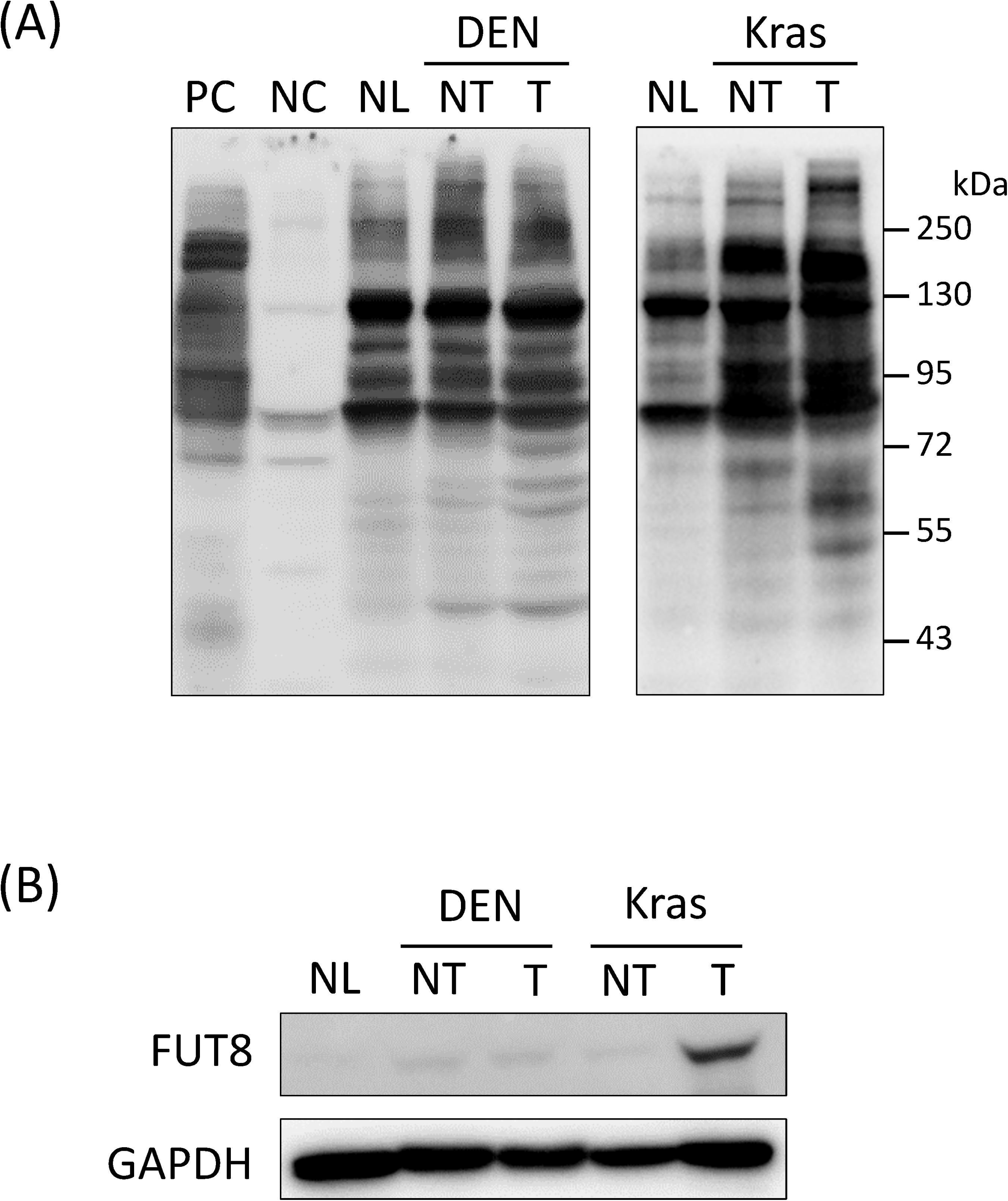
LRP1 deficiency suppresses proliferation and alters morphology in HepG2 cells (A) Western blot analysis of LRP1 in wild-type (WT) and LRP1-knockout (KO) HepG2 cells; β-actin was used as a loading control. (B) Phase-contrast micrographs of WT and LRP1 KO HepG2 cells showing representative fields (two images per condition); scale bar, 100 µm. (C) Cell proliferation of WT and LRP1 KO HepG2 cells assessed by ATP-based assays at 1, 3, and 5 days in culture. Proliferation is expressed relative to the value at day 1 for each cell line (set to 1.0); n = 6 per group, mean ± SD, * p < 0.01 (Student’s t-test with Bonferroni correction). (D) Western blot analysis of EGFR in wild-type (WT) and LRP1-knockout (KO) HepG2 cells; β-actin was used as a loading control. (E) Semi-quantitative RT-PCR analysis of *EGFR* mRNA expression in HepG2 WT and LRP1 KO cells. Expression levels were normalized to *ACTB* and are shown relative to WT cells (set to 1.0); n = 3 per group, mean ± SE, * p < 0.05 (Student’s t-test).

## Discussion

In the Kras^G12D^ mouse model, fucosylation was enhanced in liver tumor tissues compared with the surrounding non-tumor liver tissue. FUT8 protein levels were increased in mouse HCC, consistent with previous reports linking FUT8 upregulation to HCC progression(2, 4, 5). Glycoproteomic analysis of AAL-enriched fractions from Kras tumors further identified Low-density lipoprotein receptor-related protein 1 (LRP1) as a major fucosylated protein.

LRP1 is a multifunctional endocytic and signaling receptor that binds a wide range of ligands, including lipoproteins, protease–inhibitor complexes, and extracellular matrix proteins(27). LRP1 has been reported to modulate diverse processes relevant to cancer biology, such as proliferation, migration, invasion, and responses to growth factors, and its expression level has been associated with clinical outcome in several malignancies(28). Intriguingly, LRP1 can exhibit either tumor-promoting(29, 30) or tumor-suppressive(31, 32) effects depending on the tumor type and microenvironmental context. Furthermore, previous work in Fut8-deficient mice demonstrated that loss of core fucosylation impairs LRP1-mediated ligand clearance, indicating that core fucose is crucial for full LRP1 function *in vivo*(33). These observations raise the possibility that FUT8-dependent core fucosylation may fine-tune LRP1 activity in the liver and thereby influence hepatocarcinogenesis. In our experiments with immortalized mouse hepatocytes, both *Fut8* genetic deletion and pharmacological inhibition resulted in increased *Lrp1* mRNA expression and marked accumulation of LRP1 protein, which contrasts with the upregulation of FUT8 and LRP1 in Kras mouse liver tumors. This discrepancy suggests that FUT8 can influence LRP1 at two distinct levels: directly, by modifying its *N*-glycans and thereby altering receptor function, and indirectly, by affecting transcription and/or stability of LRP1 in a context dependent manner.

LRP1-deficient HepG2 cells showed reduced proliferation and a marked morphological change, forming compact, rounded colonies instead of polygonal morphology. In addition, LRP1 knockout decreased total EGFR protein. Consistent with our observations, a recent study reported that LRP1 suppression reduces EGFR and inhibits proliferation in gastrointestinal cancers, supporting the idea that LRP1 positively regulates EGFR-linked growth pathways(34). Moreover, in several non-hepatic cancers, high LRP1 expression has been associated with increased invasiveness or poor clinical outcome(35), suggesting that LRP1 can exert pro-tumor effects. At the same time, however, LRP1 has also been reported to exert tumor-suppressive effects in certain *in vivo* HCC settings, where low LRP1 expression is associated with malignant progression and poor prognosis(36, 37). These apparently conflicting observations indicate that the impact of LRP1 on EGFR and proliferation is highly context-dependent. In particular, differences in tumor microenvironment, cellular background, and glycosylation status —most notably core fucosylation—are likely to influence how LRP1 shapes signaling in HCC. In this study, only a single LRP1-knockout HepG2 clone survived antibiotic selection and showed markedly reduced proliferation, which is consistent with an important role for LRP1 in sustaining growth of HepG2 cells under standard culture conditions, although clonal selection effects cannot be excluded. Importantly, LRP1 knockout decreased total EGFR protein and was associated with an accumulation of cells in the G2/M phase, suggesting that LRP1 can contribute to EGFR-linked cell-cycle progression in HepG2 cells. While EGFR inhibition is often linked to G1 arrest, the cell-cycle response can vary by cellular context, and G2/M arrest has also been described(38, 39). In the present setting, however, baseline propidium iodide uptake was altered in the LRP1-knockout clone, which limits the precision of the cell-cycle analysis, so we interpret the apparent G2/M accumulation mainly as supportive of the reduced proliferation phenotype rather than as definitive evidence for a specific arrest pattern.

Our data also indicate that the regulation of LRP1 fucosylation differs between hepatocarcinogenesis models. DEN-induced tumors, which arise in the setting of chronic liver injury and fibrosis(40, 41), did not show a global increase in AAL-reactive fucosylation or FUT8 upregulation. By contrast, Kras-driven liver tumors developing under strong oncogenic Ras signaling exhibited a marked increase in AAL lectin binding and LRP1 protein levels. These observations suggest that the contribution of fucosylation, and possibly LRP1 fucosylation, to hepatocarcinogenesis is context-dependent and may be more pronounced in oncogene-driven tumors than in chemically induced tumors, although the underlying mechanisms remain to be clarified.

Collectively, our findings suggest that the FUT8–LRP1 axis can support growth-related signaling in liver cancer, at least in the specific models examined. It also highlights LRP1 as a glyco-regulated receptor that should be considered when targeting FUT8 and core fucosylation in HCC.

## Acknowledgement

Current affiliation of Y.M. is Division of Chemistry, Department of Materials Engineering Science, Graduate School of Engineering Science, The University of Osaka.

The authors acknowledge the use of an artificial intelligence–based assistant (Perplexity, powered by GPT) for English language editing and refinement of the manuscript text during March–April 2026. The AI was not used for data generation, data analysis, or drawing scientific conclusions, and all interpretations and final manuscript contents were reviewed and approved by the authors.

## Author Contributions

J.K., Y.K., and E.M. conceived and designed the experiments; M.U and K.O. performed the experiments; A.O., M.U., K.O and S.T. analyzed the data; Y.M. and K.F. contributed reagents; Y.M. and H.H., and Y.O. contributed materials; J.K. and A.O., and E.M. wrote the paper. All authors reviewed, edited and approved the final manuscript.

