## Supplementary Fig S1 for "FUT8-mediated core fucosylation modulates growth-related functions of LRP1 in liver cancer cells"

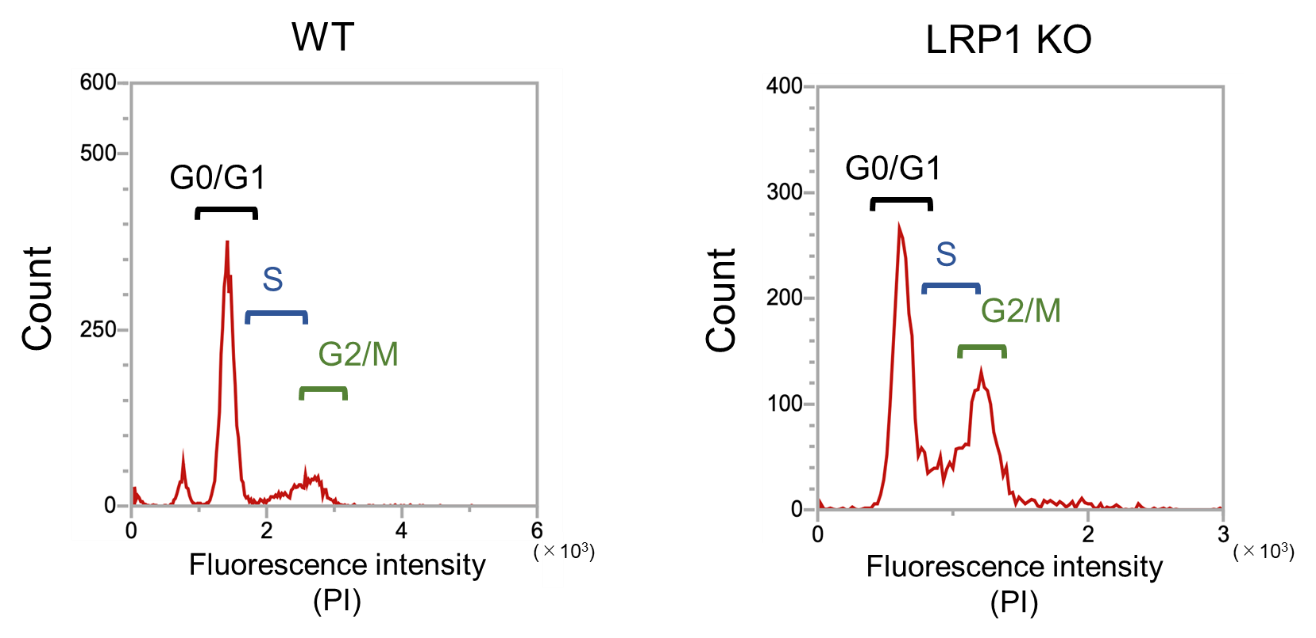

**Supplementary Fig S1.**

**Cell‑cycle distribution in HepG2 WT and LRP1 KO cells**

Cell‑cycle analysis of HepG2 WT and LRP1 KO cells by flow cytometry, showing the distribution of cells in the G1, S, and G2/M phases.
